# Limited thermal plasticity and geographic divergence in the ovipositor of *Drosophila suzukii*

**DOI:** 10.1101/670356

**Authors:** Ceferino Varón-González, Antoine Fraimout, Arnaud Delapré, Vincent Debat, Raphaël Cornette

## Abstract

Phenotypic plasticity has been repeatedly suggested to facilitate adaptation to new environmental conditions, as in invasions. Here we investigate this possibility by focusing on the worldwide invasion of *Drosophila suzukii*: an invasive species that has rapidly colonized all continents over the last decade. This species is characterized by a highly developed ovipositor, allowing females to lay eggs through the skin of ripe fruits. Using a novel approach based on the combined use of SEM and photogrammetry, we quantified the ovipositor size and 3D shape, contrasting invasive and native populations raised at three different developmental temperatures. We found a small but significant effect of temperature and geographic origin on the ovipositor shape, showing the occurrence of both geographic differentiation and plasticity to temperature. The shape reaction norms are in turn strikingly similar among populations, suggesting very little difference in shape plasticity among invasive and native populations, and therefore rejecting the hypothesis of a particular role for plasticity of the ovipositor in the invasion success. Overall, the ovipositor shape seems to be a fairly robust trait, indicative of stabilizing selection. The large performance spectrum rather than the flexibility of the ovipositor would thus contribute to the success of *D. suzukii* worldwide invasion.

## 1. Introduction

Phenotypic plasticity is a pervasive feature in nature (West-Eberhard, 1989) and a major response to changing environmental conditions (Bradshaw, 1965). Because it may facilitate the colonization of new environments (e. g. Lande, 2015), it has been suggested that plasticity may play an important role in biological invasions: accordingly, invasive populations are expected to be more plastic than non invasive populations (Davidson et al., 2011; Lande, 2015; Lee and Gelembiuk, 2008; Richards et al., 2006; Yeh and Price, 2004). Although often discussed theoretically (Chevin et al., 2010; Via and Lande, 1985), this hypothesis has been comparatively rarely tested (Richards et al., 2006), in particular in animal species (Fraimout et al., 2018; Loh et al., 2008).

*Drosophila suzukii* has received much attention over the last 10 years, as it has colonized multiple countries worldwide (Fraimout et al., 2017) and induced severe losses in agriculture (Asplen et al., 2015; Farnsworth et al., 2017; Mazzi et al., 2017). This species has been extensively collected to test hypotheses about the role of plasticity during its invasion (e.g. Clemente et al., 2018; Fraimout et al., 2018; Poyet et al., 2015; Shearer et al., 2016). However, plasticity largely depends on the environmental factor considered and the morphological trait under study (Fraimout et al., 2018; Nijhout and German, 2012; Nijhout et al., 2014; Shingleton et al., 2009). For *D. suzukii*, temperature has been frequently chosen as the factor inducing phenotypic plasticity due to its pervasive effect on insect development (e. g. Atkinson, 1994; Crill et al., 1996; David et al., 1997) and its importance in shaping the distribution of Drosophila species (David et al., 1997). Different morphological structures such as wings, thorax and ovipositor have been investigated (e. g. Clemente et al., 2018; Fraimout et al., 2018; Shearer et al., 2016). The ovipositor is a particularly interesting structure owing to the reproductive behavior of this species: *D. suzukii*’s damaging potential is indeed due to its over-developed ovipositor, used to pierce through the skin of ripening fruits and lay its eggs (Atallah et al., 2014). It is well-known that fruits texture is strongly affected by temperature (e. g. Bourne, 1982): specifically, their firmness and resistance to puncture tends to decrease with increasing temperature (e. g. Khazaei and Mann, 2004). It is thus conceivable that *D. suzukii* ovipositor might present some adaptive plasticity to temperature, allowing it to pierce fruits skins of (thermally induced) varying resistance. An alternative hypothesis is that it might rather be under stabilizing selection, as has been suggested in *D. melanogaster* for genitalia (Shingleton et al., 2009), in which case we should expect a reduced sensitivity to temperature.

The ovipositor is a microscopic 3D structure (about 500 μm). 3D characterization of its shape is essential to recover all the possible features involved in its performance and therefore to link its morphology to the possible selective forces affecting it. 2D approximations of 3D structures might be troublesome because all the variation recovered by one physical dimension would be missing and that might affect the analysis (Buser et al., 2018; Cardini, 2014). Finally, the complete description of shape may be particularly important for assessing the ovipositor plasticity: a 2D analysis could lead to underestimations of the plastic shape change when the plastic variation is not recovered among the shape descriptors. We thus developed an approach based on the combination of Scanning Electron Microscopy (SEM)-based photogrammetry and 3D geometric morphometrics allowing to finely depict and quantify the ovipositor 3D shape and its variation.

In this study we analyze the plastic response of the ovipositor shape to developmental temperature in three different geographic populations of *D. suzukii*, including a population from the native range (Japan) and two populations from the invaded range (France and USA). These three geographic populations represent the three most genetically divergent populations of the distribution (Fraimout et al., 2017). By contrasting laboratory lines derived from native and invasive populations, we (1) investigate whether there is any genetic divergence in the ovipositor shape across the distribution range; (2) quantify the ovipositor plasticity to temperature and (3) investigate whether plasticity is higher in invasive populations, as predicted if plasticity played a role in the invasion success, possibly allowing *D. suzukii* to exploit a larger diversity of substrates in varying thermal conditions.

## 2. Materials and methods

### (a) Samples

Adult flies were sampled in 2014 using banana bait traps and net swiping in three different regions: one belonging to the native range (Sapporo, Hokkaido, Japan) and two to the invasive range (Paris, France and Dayton, Oregon, USA). Ten isofemale lines per locality were stocked so that they performed single matings separately and the F1 offspring was expanded in consecutive series of vials (Hoffmann and Parsons, 1988). These stocks were maintained at 22°C on a medium with corn starch, yeast with antibiotics and hydroxyl-4 benzoate. Female flies were left to oviposit for 24 hours in two separate sets of 20 vials and after oviposition was checked parent flies were removed. Then, two batches were placed in two incubators: one set of eggs was stored at 16°C and other one at 28°C (keeping a third at 22°C). Therefore, for each population and temperature we produced ten isofemale lines in separate rearing vials with single matings at three different experimental temperatures: i.e. 30 lines per geographic population. The position of the incubators was assigned randomly and they were kept at the experimental temperatures until 2 days after the emergence. These experiments were originally conducted to run the analyses published in Fraimout et al. (2018).

Final samples consisted on 20 individuals from Paris raised at 16°C, 11 at 22°C and 13 at 28°C. 19 individuals from Sapporo raised at 16°C, 20 at 22°C and 23 at 28°C and 14 individuals from Dayton at 16°C, 6 at 22°C and 13 at 28°C.

### (b) Electronic microscopy

For each fly, the ovipositor was detached from the body – the two valves being kept in connection – and the connective tissues were manually removed. Because all the specimens were conserved in alcohol, no deformation was produced during the removal of the ovipositors. Then, they were photographed using an environmental scanning electron microscope (ESEM). Images were collected in low vacuum (0.37 Torr) with a large field low vacuum SED detector (LFD) using a FEI Quanta 200 FEG operating at 15 Kv at a working distance of 10 mm.

From each ovipositor 52 pictures were taken describing two semicircular trajectories, perpendicular between them. That allowed recovering information from all different angles of each specimen.

### (c) Photogrammetric reconstruction

The 3D reconstruction of each ovipositor was inferred using photogrammetry (Figure 1), the technique allowing the 3D representation of an object from a set of pictures. The photogrammetric process starts with the alignment of the pictures obtained from the ESEM, i. e. the recognition of analogous parts among pictures. Where difficulties for the picture alignment were found, a mask was applied to select just the ovipositor within the pictures and discard the background, facilitating the correct alignment of the pictures. The inference of the distances among analogous pixels allow the inference of the position of these pixels in a 3D space (i. e. the transformation of pixels in voxels). Once this first point cloud was inferred, all the voxels not corresponding to the ovipositor itself were removed. This cleaning fastens the next step, the reexamination of the picture alignment once a first point cloud was build in order to obtain more analogous voxels. As a result, from the first point cloud we obtained a dense cloud. Finally, a mesh was built based on the dense cloud with no *a priori* about the final shape (arbitrary surface type). All reconstructions were done in PhotoScan (Agisoft, 2014).

**Figure 1.**
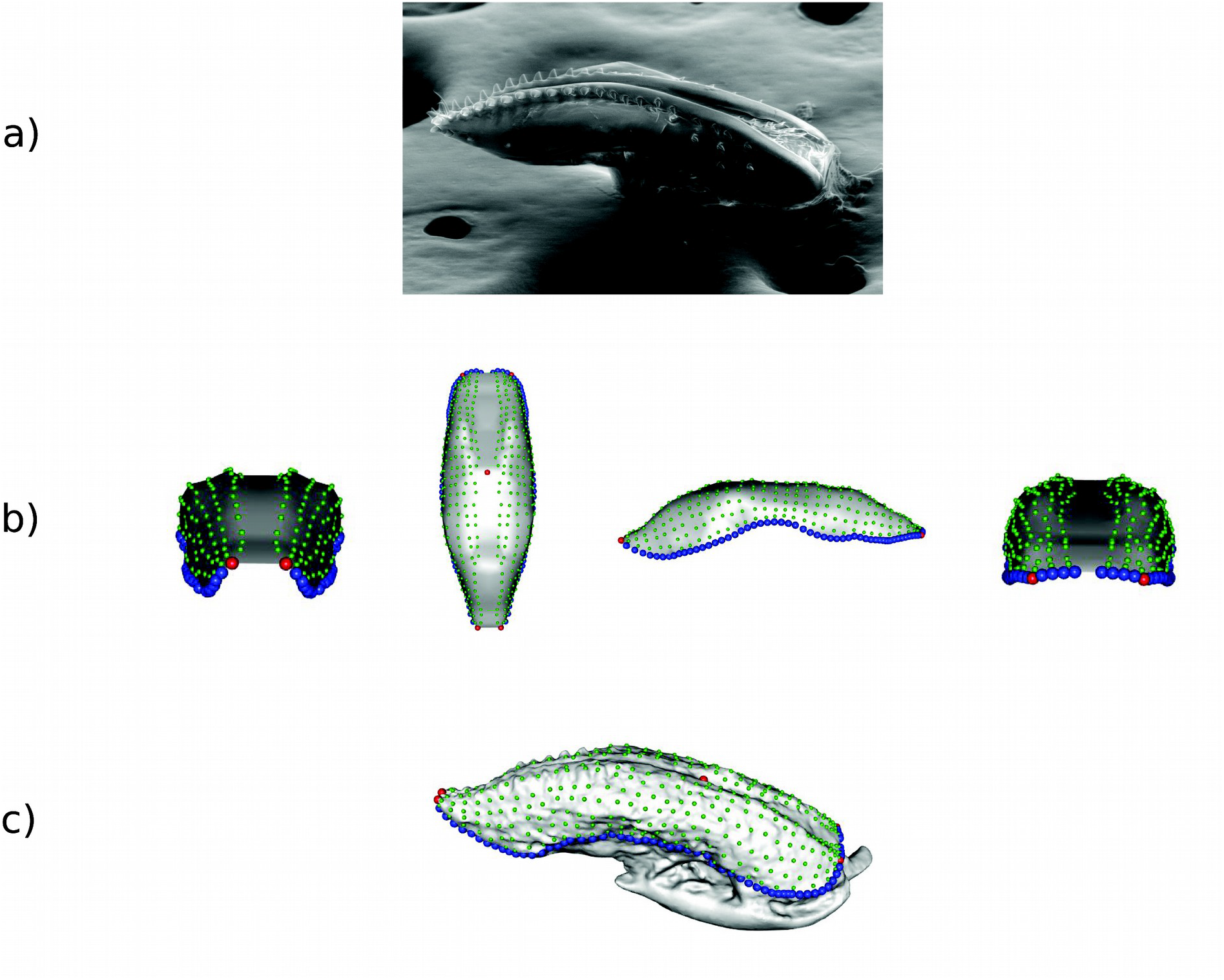
Ovipositor at electronic microscopy (a), template (b) and ovipositor phenotyping (c). Once the ovipositor pictures at electronic microscopy were obtained (a) and the 3D reconstruction of the ovipositor was done, we build a template with a simplified shape of an ovipositor (b) where we placed landmarks (red), semilandmarks (blue) and surface semilandmarks (green). This template was then projected to each 3D reconstruction to obtain the 3D landmarks characterizing the ovipositor shape (c).

Because many of the reconstructions were built using a mask, the scale bar present in the pictures did not appear in the 3D reconstructions and therefore we could not give the correct scale of each 3D model during the reconstruction process. For that, once the 3D models were obtained we measured the real lengths of the ovipositors in the pictures using ImageJ 1.51j8 (Rasband, 2012) and then we scale each ovipositor in MeshLab v2016.12 (Cignoni et al., 2008). The advantage of MeshLab is that the linear measurements of the object do not consider its surface curvature (i.e. it uses Euclidean distances), the same as picture measurements. In any case, to avoid any possible deformation due to the picture perspective we used the dorsal pictures of the ovipositor and the dorsal 3D view of the ovipositor (the flattest part).

### (d) Morphometric analyses

A set of 5 landmarks and three curves containing in total 130 semi-landmarks were defined in each 3D mesh (Figure 1) (Gunz and Mitteroecker, 2013). One pair of landmarks was fixed at the most distal part of the ovipositor and other pair at the most proximal part. The fifth fixed landmark was placed on the dorsal area, at the ovipositor opening. Two curves with 60 semilandmarks each were placed on the ovipositor sides. The other 10 semilandmarks surround the proximal area of the ovipositor. Landmarking was performed on Landmark Software (Wiley et al., 2005). Then, we created a template replicating a simplified form of an ovipositor (Figure 1), composed of 394 surface points. Landmarks, semilandmarks and 394 surface semilandmarks were digitized on the template and they were used to deform the template via thin plate spline. Finally, all landmarks were projected on the ovipositor and they slided to minimize bending energy taking into account the ovipositor object symmetry (Gunz et al., 2005). In total 529 landmarks described the ovipositor shape for each individual. This process follows the protocol described by Botton-Divet et al. (2015). The template was created with Meshlab (Cignoni et al., 2008) and the position of these landmarks and the subsequent sliding were performed with the R package Morpho (Schlager, 2017).

To assess the quality of the 3D shape reconstruction we replicated the reconstruction process five times on two individuals from the same geographic population and raised at the same temperature (two Sapporo individuals raised at 16°), so the variance between individuals was minimized as much as possible. The reconstructions was done on each one of these two individuals five times and the landmarks were collected on each of the ten meshes. A multivariate model was run with the function *procD.lm* (Adams et al., 2018) to test for the amount of variance explained by inter-individual variation in relation to the variation explained by the reconstruction and landmarking processes (residuals).

Differences among populations and temperatures were explored using a between-group PCA (Mitteroecker and Bookstein, 2011). A permutation test was run this transformation of the space was computed to test the significance of the differences among populations. Individuals from all groups (populations and temperature) were randomly shuffled and new pairwise Procrustes distances among group means computed. 10000 permutations were run and significance levels obtained as the proportion of Procrustes distances less extreme. Because the permutations are run on the whole sample and significance tests are not independent, no correction for multiple comparisons is needed. The effect on shape of the temperature and population factors as well as their interaction were tested with a linear multivariate model and permutation tests as performed in the geomorph function *procD.lm* (Adams et al., 2018). The effect size for each factor was assessed by Z, an estimator based on the F-statistic (Collyer et al., 2015). The effect the two factors on the centroid size was assessed with a two-way ANOVA.

To further compare the plastic responses among populations we used the trajectory analysis method developed by Collyer and Adams (2013). This approach specifically tests the similarity between trajectories depicting shape changes in the multivariate shape space and it can be readily transposed to the analysis of shape reaction norms. With this analysis, three different aspects of the shape change are studied: the amount of shape change as the trajectory path length (size), the pattern of shape covariation as the difference in angles among the first principal component of each trajectory (direction) and the differences in trajectory shapes (shape) as Procrustes distances between pairs of phenotypic trajectories. Although these three aspects of plasticity are somehow related, they do not look at the same effects temperature may have on shape variation. The amount of shape change reflects whether the effect of temperature on shape variation (i. e. plasticity) is larger or smaller in some populations. The pattern of shape covariation whether temperature affects the correlation patterns among the different landmarks, changing which landmarks covary together in response to temperature and in the degree and sign of these covariations. Last, the differences in trajectory shapes study the existence of differences among shapes at the same temperature. The statistical assessment of these three features is based on the simulated resampling from a distribution characterized by the difference in path length, angle or distances respectively and their standard deviations.

Allometry was quantified using a linear model of the logarithm of the centroid size against symmetric shape (Monteiro, 1999). A general allometric pattern was expected given the pervasive effect of temperature on size in insects (David et al., 1997) as well as previously published effects in 2D (Clemente et al., 2018). Differences in the allometric slopes among geographic populations were also assessed. Because the allometric patterns are expected to be primarily influenced by temperature variation, we would expect the differences in allometric slopes and the differences in reaction norms to be analogous. Differences among slopes were tested with an ANOVA. All morphometric tests were applied in the R package geomorph (Adams et al., 2018).

Finally, we investigated the degree of relative robustness of the ovipositor, by comparing its variation with that of the wing, as assessed on the same samples by Fraimout et al. (2018). For size, we simply computed the coefficient of variation both within and among temperatures (i. e. among mean centroid sizes per temperature), for both structures. To test for differences in the coefficient of variation of size between structures we used the modified signed-likelihood ratio test, as computed in the R package *cvequality* (Krishnamoorthy and Lee, 2014). Comparing the shape variability of two different objects is challenging, because they lie in different shape spaces and no direct multivariate extension of the coefficient of variation can be applied. We used Mahalanobis distances among temperatures, computed independently for the two structures, as a measure of their relative sensitivity to temperature. Because this distance measures the difference between groups relative to the within group variation (Klingenberg and Monteiro, 2005), it could be comparable between structures with a note of caution. Because our hypothesis is that the ovipositor will show more robustness, it may also show lower within-group variation and that would increase distances. In this particular case, this feature makes our estimates more conservative but further applications of this method would require appropriate justification. As distance measures are affected by the data dimensionality, we estimated the Mahalanobis distances on the same number of principal components for each dataset (26 principal components: 100% of the fly shape variation and 96.94% of the ovipositor shape variation). The statistical assessment of the difference in plasticity between temperatures was done with an *ad-hoc* permutation test under the null hypothesis that the difference in plasticity between structures is equal to zero. In it, for the ovipositor and the wing independently, all distances from individuals at one temperature to individuals at other temperature were estimated. Then, all within-structure among-temperature distances were randomly assigned to two groups and the difference between the means of these two groups stored. The permutation was run 1000 times and a distribution of differences in plasticity (centered in zero) was generated. The empirical differences were considered significant if their real value fell out of the 95% of the values in the distribution. To obtain the distances among temperatures we applied the function *CVA* in the R package Morpho (Schlager, 2017). All analyses and data management were conducted in RStudio version 1.1.442 (RStudio Team, 2016).

## 3. Results

The 3D shape reconstruction of the ovipositors allowed us to assess the ovipositor 3D shape variation precisely. We found a significant effect of temperature and geographic variation on the ovipositor size and 3D shape, but the effects appeared weak and all nine experimental groups were not fully discriminated (Table 1). Although the interaction between geographic and temperature factors was significant in the multivariate model, no differences among shape trajectories or allometric slopes in response to temperature were detected among geographic populations (Figure 2).

**Table 1.** Discriminant analysis for temperature and geographic factors. 1000 permutations using Procrustes distances between group means were run with the function *groupPCA* of the R package Morpho. No significant results are shaded.

|  | Paris 16° | Paris 22° | Paris 28° | Sapporo<br>16° | Sapporo<br>22° | Sapporo<br>28° | Dayton<br>16° | Dayton<br>22° |
| --- | --- | --- | --- | --- | --- | --- | --- | --- |
| Paris 22° | 0.0058 |  |  |  |  |  |  |  |
| Paris 28° | 0.0082 | 0.0104 |  |  |  |  |  |  |
| Sapporo<br>16° | 0.1224 | 0.0034 | 0.0004 |  |  |  |  |  |
| Sapporo<br>22° | 0.0002 | 0.0063 | 0.0001 | 0.0036 |  |  |  |  |
| Sapporo<br>28° | 0.0107 | 0.0001 | 0.0055 | 0.0550 | 0.0053 |  |  |  |
| Dayton<br>16° | 0.0306 | 0.0115 | 0.0001 | 0.0415 | 0.0002 | 0.0002 |  |  |
| Dayton<br>22° | 0.0107 | 0.3861 | 0.0010 | 0.0790 | 0.0208 | 0.0007 | 0.2093 |  |
| Dayton<br>28° | 0.0460 | 0.0044 | 0.0060 | 0.1402 | 0.0193 | 0.2199 | 0.0136 | 0.0254 |

**Figure 2.**
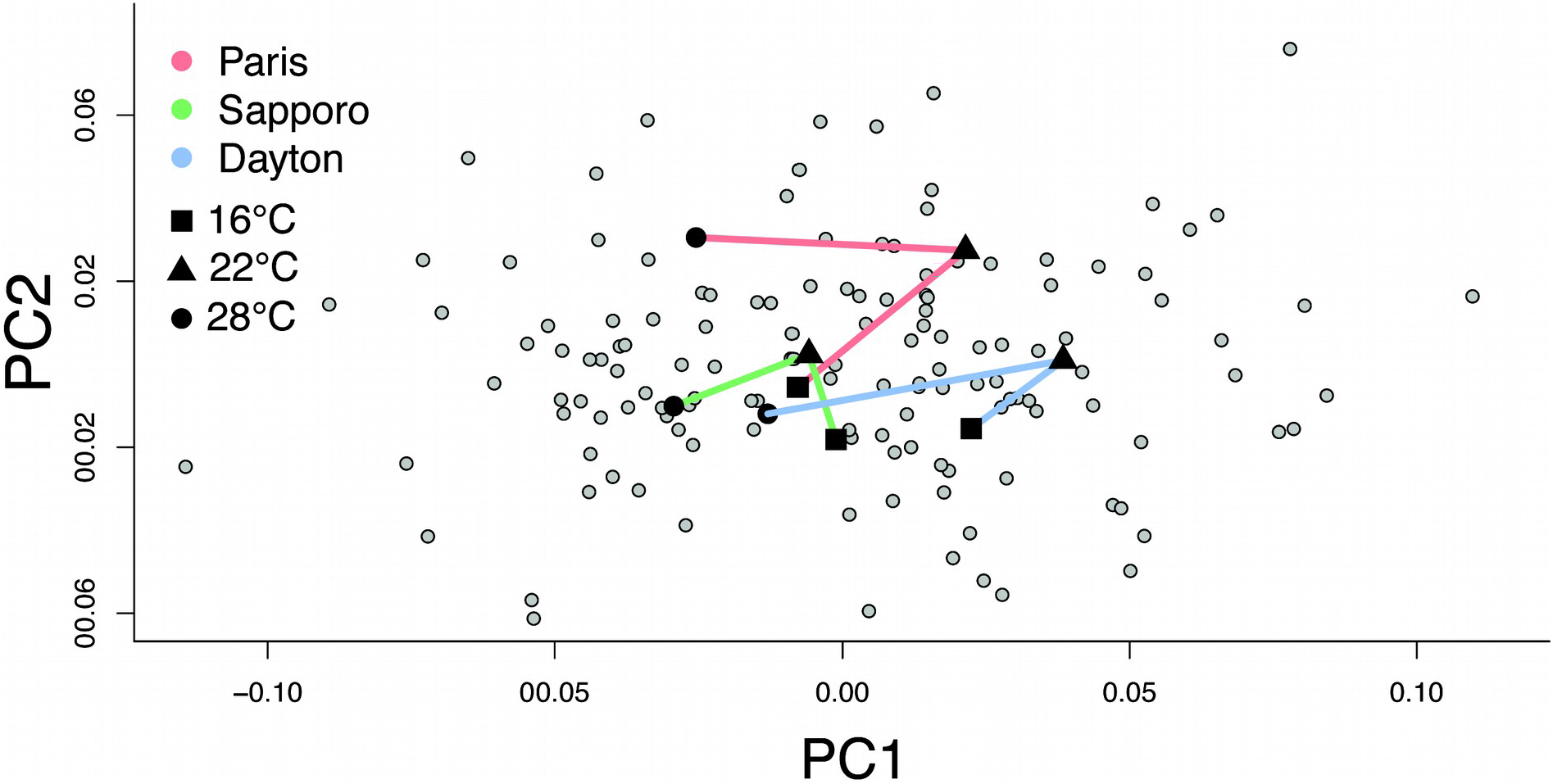
Ovipositor 3D shape variability and plasticity trajectories in response to developmental temperature. First two principal components of the ovipositor shape for individuals (gray) and temperature means for each geographic population (black; square: 16°C, triangle: 22°C, circle: 28°C). The three temperature levels for each geographic population are joined so the reaction norms can be visualized for each population (Paris: red, Sapporo: green, Dayton: blue). We can observe the overlap among reaction norms and the similarity in their trajectories, suggesting similar plasticities among populations.

### (a) Measurement error

The repeated reconstruction of the 3D shape of the two individuals from Sapporo raised at 16°C showed that the variation in the reconstruction process was almost four times smaller than variation between individuals (MS_IND_/MS_RES_ = 3.92, *p* = 0.011). Although substantial, measurement error due to 3D reconstruction and landmarking processes should not preclude detection of genuine variation among individual ovipositors.

### (b) Temperature and population effects

Overall, both temperature (Z = 5.27, *p* < 0.001) and geography (Z = 4.72, *p* <0.001) had a significant effect on ovipositor shape. In addition, temperature interacted with geography in their association with shape (Z = 1.95, *p* = 0.026), suggesting a different effect of temperature among geographic populations. The pairwise comparisons between geographic samples showed that the significance of this interaction was driven by a subtle difference between Sapporo and Paris populations (Z = 1.96, *p* = 0.035).

The temperature shift from 22° to 16°C is associated with a narrowed ovipositor overall (Figure 3). At 16° the ovipositor seemed to be elongated and flatter, producing an inner folding of the lateral parts of the ovipositor within the structure and therefore smaller and plane lateral parts. The increase from 22° to 28°C produced again an overall narrowing of the ovipositor (although less pronounced than at 16°) and the widening of the anterior part of the ovipositor. In comparison to Sapporo population, Paris population showed a narrower posterior part and more folding on the lateral parts, which were smaller. Dayton seemed the most elongated geographic population and the one with the narrowest anterior part.

**Figure 3.**
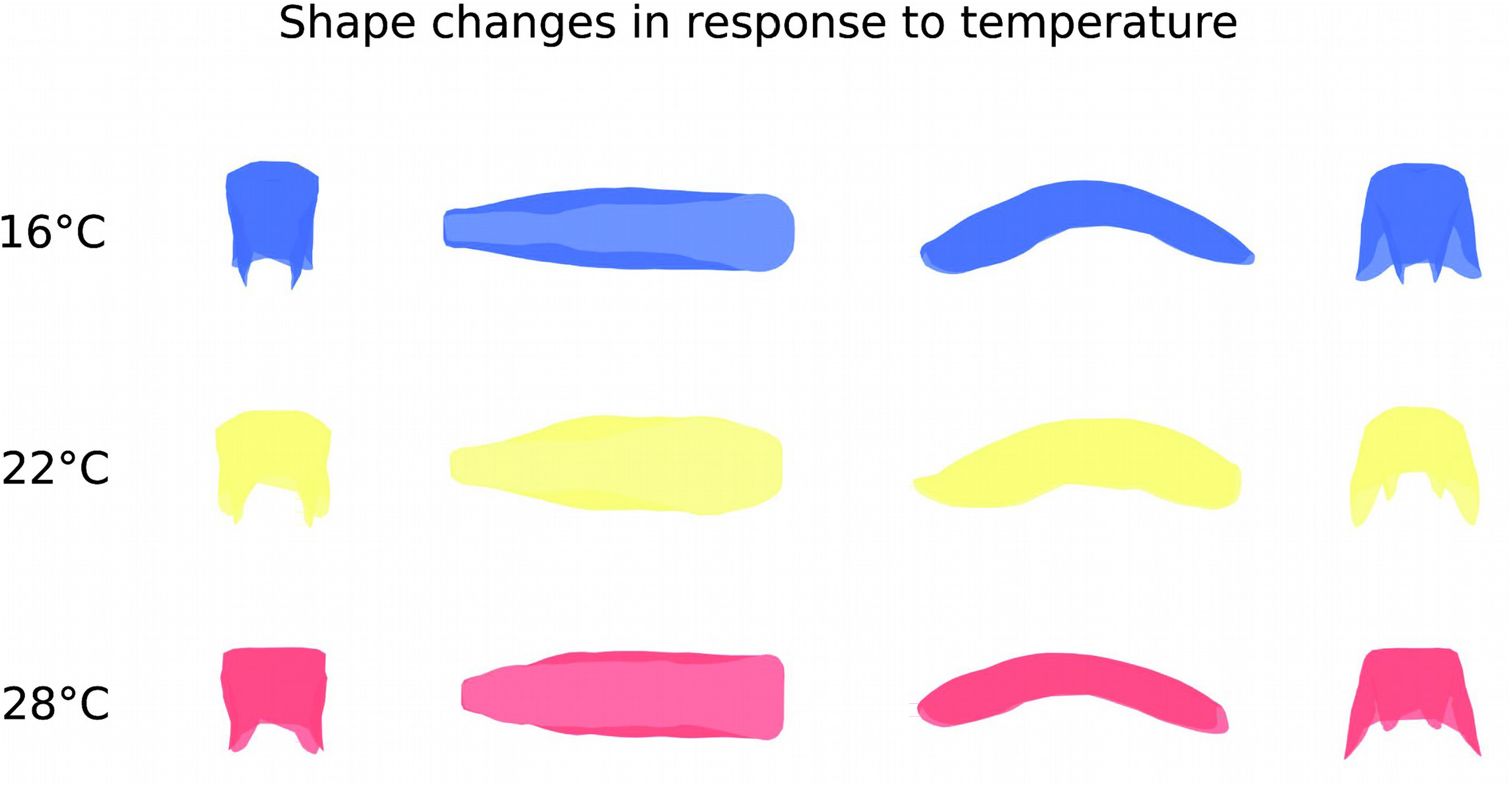
Effect of developmental temperature on the ovipositor 3D shape. While the ovipositor shape at 22°C (center row) represents the approximate real shape of the three populations at that temperature, morphologies at extreme temperatures are represented as exaggerated versions (five standard deviations) of the linear transformation from 22°C to each temperature. Therefore, the linear transformation from 16° to 28°, not biologically meaningful as the effect of temperature is not linear, is not represented. 3D shapes are captured by four different perspectives (from left to right: posterior, dorsal, lateral and anterior). Shape changes were obtained with the Morpho library. See results for the description of the shape changes.

**Figure 4.**
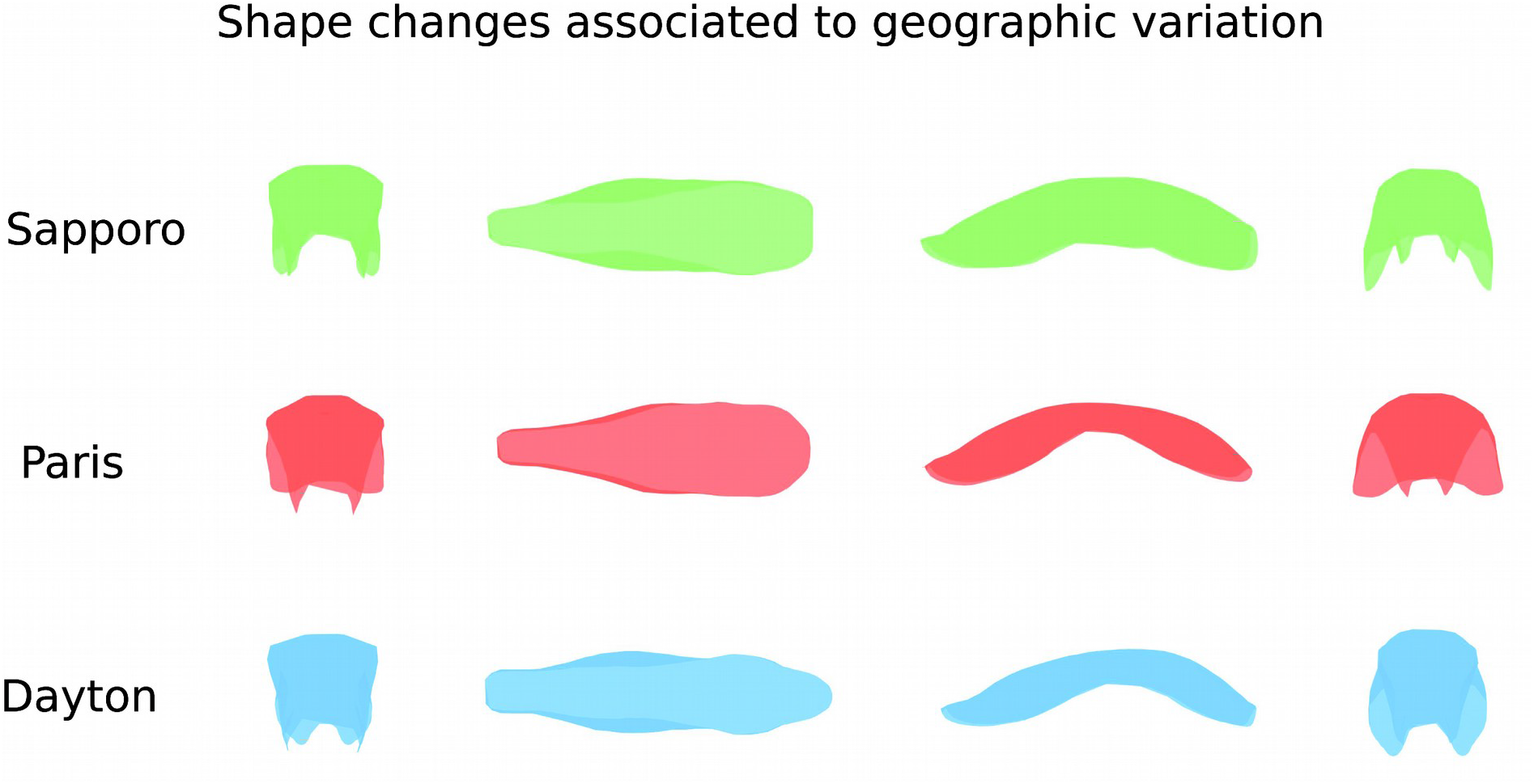
Effect of geographic variation on the ovipositor 3D shape. Here we represent the exaggerated linear deformation (three standard deviations) from the overall mean shape to each geographic population shape. 3D shapes are captured by four different perspectives (from left to right: distal, dorsal, lateral and proximal). Similarly to Figure 3, the lineal transformation from the Paris to Dayton population does not make biological sense since both come from a Japanese population (Fraimout et al., 2017). Therefore, Sapporo population is represented by its true mean shape and the other two population as a linear transformation from the former to each of the latter populations. Shape changes were obtained with the Morpho library. See results for the description of the shape changes.

The trajectory analysis showed a striking conservation of the shape variation patterns among geographic populations (Figure 2). Trajectories for all geographic populations showed very similar path lengths (Paris = 0.10, Sapporo = 0.08, Dayton = 0.10) and no difference was detected (Sapporo-Paris: effect size = −0.02, *p* = 0.41, Sapporo-Dayton: effect size = −0.48, *p* = 0.63, Paris-Dayton: effect size = −1.05, *p* = 0.87). Although angles among populations showed larger variation, no difference among trajectory angles was found (Sapporo-Paris: angle = 120.56°, Effect size = 0.98, *p* = 0.977, Sapporo-Dayton: angle = 100.36°, Effect size = 0.48, *p* = 0.361, Dayton-Paris: angle = 41.92°, Effect size =−0.96, *p* = 0.77). Similarly, shape differences among trajectories were no significant (Sapporo-Paris: Procrustes distance = 0.10, effect size = −1.16, *p* = 0.89, Sapporo-Dayton: Procrustes distance = 0.25, effect size = −0.04, *p* = 0.47, Paris-Dayton: Procrustes distance = 0.17, effect size = −0.70, *p* = 0.74).

### (c) Size variation and allometry

The ovipositor size was found to decrease with increasing temperature (Figure 5, F_2, 130_ = 92.31, *p* < 0.001). Geography also showed a significant effect on the ovipositor size (Figure 5, F_2, 130_ = 14.875, *p* < 0.001), Dayton populations being larger than Paris. No interaction between temperature and population effects was detected (F_4, 130_ = 2.138, *p* = 0.08), suggesting that the plasticity of ovipositor size was conserved across populations.

**Figure 5.**
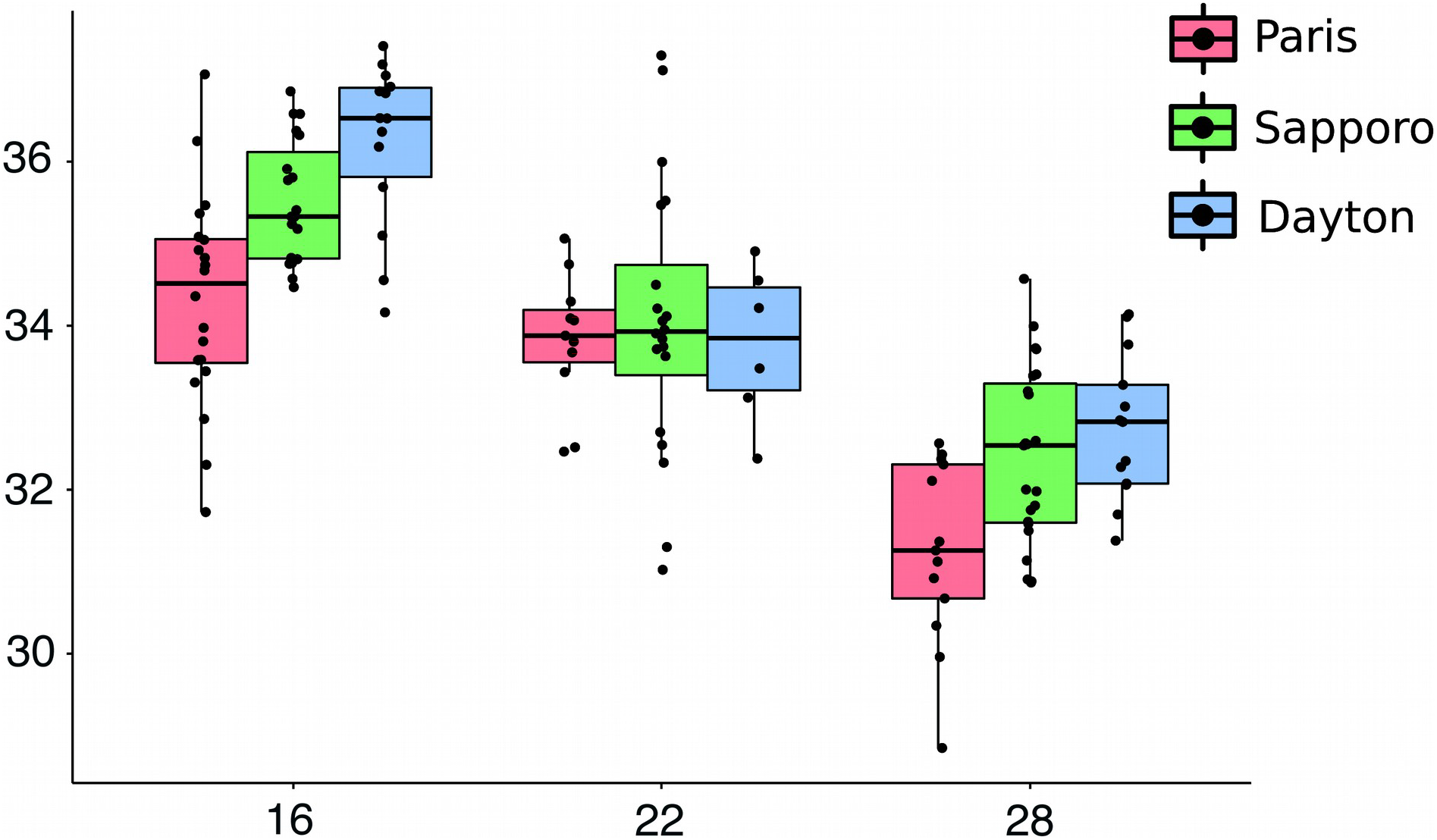
Effect of developmental temperature on the ovipositor centroid size. Ovipositor centroid size variation in response to developmental temperature (16°C: left block, 22°C: middle block, 28°C: right block), for each population (Paris: red, Sapporo: green, Dayton: blue).

Ovipositor shape and size were correlated, so the plastic response to temperature produced a general allometric pattern (Z = 3.79, *p* < 0.001). When the allometric slope among geographic populations was compared, no significant difference was found (Z = 0.49, *p* = 0.325).

### (d) Comparison with the wing

Wing shape showed much larger Mahalanobis distances among temperatures than the ovipositor shape, suggesting that wing shape is more plastic than ovipositor shape. For the ovipositor, the distances from 22°C to the extreme temperatures are relatively stable: 2.38 to 16°C and 3.03 to 28°C. For the wing, both distances were larger but the high temperature had a stronger impact on shape: 2.87 to 16°C (*p* < 0.001) and 5.60 to 28°C (*p* < 0.001). When we look at the distances between the extreme temperatures the difference between structures became more evident: we obtained a measure of 3.74 from 16° to 28°C for the ovipositor and a measure of 7.78 for the wing (*p* < 0.001).

For the centroid size, within temperature CV were close to 3% for both the wing and the ovipositor (Wing: CV_16°C_ = 3.07%, CV_22°C_ = 3.86%, CV_28°C_ = 2.19%; Ovipositor: CV_16°C_ = 3.67%, CV_22°C_ = 3.86%, CV_28°C_ = 3.67%). When comparing CV between structures within each temperature, only significant differences at 28°C were found (MSLRT_16°C_ = 1.14, *p* = 0.286; MSLRT_22°C_ < 0.01, *p* = 0.954; MSLRT_28°C_ = 8.05, *p* = 0.005). The wing showed a much larger plastic response among temperatures than the ovipositor (CV_WING_ = 14.28%, CV_OVIPOSITOR_ = 4.55%, MSLRT = 65.42, *p* < 0.001).

## 4. Discussion

Our results showed significant but limited plasticity of the ovipositor shape to developmental temperature in comparison to the wing, suggesting a high robustness of the former structure against environmental variation. We also found some geographic variation associated to the ovipositor shape but its effect seemed subtle as well. This variation probably arises as a consequence of the geographic spread of this species over the last years (Fraimout et al., 2017). Although the interaction between temperature and geographic variation appeared significant, we did not find differences among reaction norms in either trajectory size, direction or shape. The allometry test confirmed these results from a different perspective: developmental temperature produces a particular relationship between the ovipositor size and shape that appeared stable among geographic populations.

Developmental temperature is a well-known factor in the origin of size and shape variation in insects (Atkinson, 1994; Ray, 1960). In the ovipositor we found the expected effect of developmental temperature (i. e. higher temperature, smaller ovipositors) (David et al., 1997) and the expected presence of allometry published for 2D analyses (Clemente et al., 2018). Our 3D approach allowed us to depict and quantify the full shape of the ovipositor and should thus allow detecting any differences among temperature and geographic factors.

In the light of our estimates, and especially if we compare the effect of temperature on the ovipositor size with that on in wing size in the same populations under the same experimental design (Fraimout et al., 2018), the ovipositor appears to be somewhat robust to temperature. This robustness is consistent with previous studies of phenotypic plasticity in Drosophila, showing a reduced variability of genitalia compared to other body parts (Shingleton et al., 2009; Shingleton et al., 2018). The mild plastic variation expressed in our experiments and the success of the invasion suggest that the ovipositor is able to perform well in a wide range of environmental conditions. One possible explanation for the limited plasticity of the ovipositor could be the effect of stabilizing selection, which could reduce its range of variation. This limited plasticity is congruent with the limited geographic variation detected and previous evidence on coevolution of the ovipositor with the male genitalia (Muto et al., 2018), expected for a trait under stabilizing selection (Eberhard, 2009; Eberhard et al., 1998). A formal Qst/FSt comparison (Ovaskainen et al., 2011) would nevertheless be necessary to test this hypothesis. Directional selection has also been showed to reduce plasticity (Bertin and Fairbairn, 2007; Bonduriansky and Day, 2003), e. g. as a consequence of the differences in the individual developmental variation in response to temperature (Dreyer et al., 2016). Because the individual is the target of selection and not particular structures within the organism, directional selection favoring certain traits may impact the population variability for a different trait. Depending on the developmental covariation between these two traits, directional selection in one trait may result in small variation for the second one.

Albeit limited, some plasticity in the ovipositor was nevertheless detected, that might have consequences on the female ability to pierce the fruits tegument. Temperature enhances fruit ripening and this change in the fruit consistency (weakening the surface) might impose new functional demands on the ovipositor morphology to successfully perforate the fruits during the oviposition (Figure 6). Although fully hypothetical, it is conceivable that the plastic shape changes reported here might have some adaptive value. This should be tested experimentally by evaluating the relative performance on a variety of substrates, of the cold and hot-generated ovipositors. Other factors like the existence of alternative selective pressures imposed on the ovipositor morphology such as sexual coevolution (Muto et al., 2018) and pleiotropic genetic effects during the ovipositor development (Green et al., 2018) might limit to such morphological adaptation.

**Figure 6.**
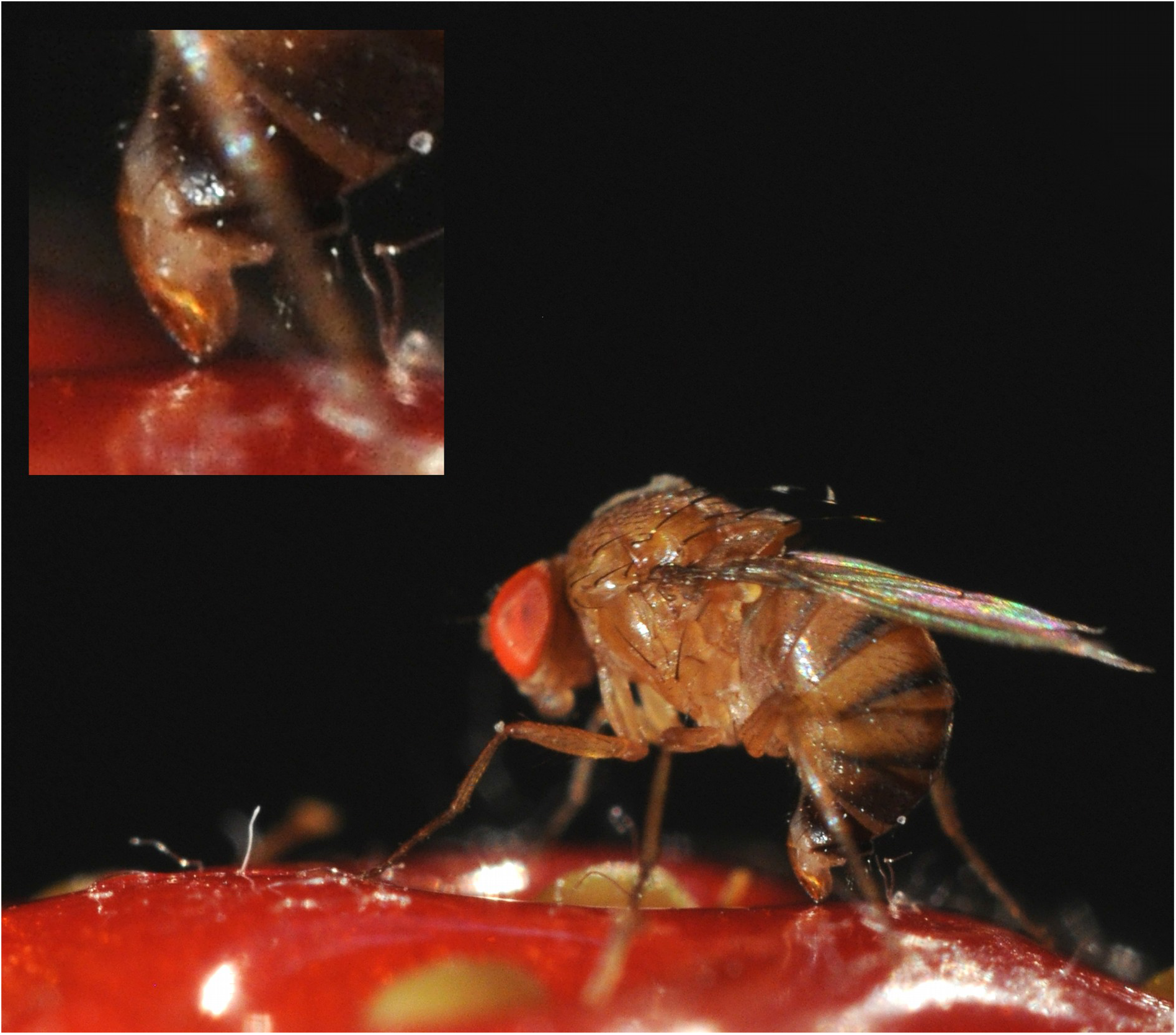
*Drosophila suzukii* **ovipositing on a strawberry**.

The lack of difference in plasticity between invasive and native populations suggests that the role of plasticity in the ovipositor during the worldwide invasion of *D. suzukii*, if any, has been limited. A similar result was found for wing shape plasticity, using males from the same populations (Fraimout et al., 2018). It has been proposed that plasticity might be transient during colonization (Lande, 2015), leaving open the possibility that plasticity might have contributed to the invasion success prior being genetically fixed. Given the speed of *D. suzukii* invasion (Fraimout et al., 2017) and the fact that all three populations show limited plastic responses, such hypothesis of ‘rapidly-evolving’ plasticity nevertheless seems unlikely.

In conclusion, while we detected some genetic divergence among populations and some thermal plasticity, phenotypic variation of the ovipositor was very limited, suggesting a high phenotypic robustness indicative of a history of stabilizing selection. The lack of difference in plasticity among populations suggests that the ovipositor large performance spectrum and phenotypic robustness rather than its plasticity would contribute to *D. suzukii* invasive success.

## Acknowledgements

We would like to thank K. Tamura, M. Toda, P. Shearer, S. Fellous and T. Schlenke for their help during the fly sampling. We also thank M. Guillaume for her help with the fly stock maintenance and F. Peronnet for the rearing medium we used. ESEM imaging was performed at the Plateform MEA, University of Montpellier. We also would like to thank Géraldine Toutirais of the “Plateau Technique de Microcopie Electronique du MNHN” and Céline Houssin for her technical expertise in electronic microscopy. S. Gerber helped with the interpretation of some of the methods. This article was also improved by the comments made during the SWING project meetings and the different people attending the seminars where it was presented, as the GDR PlasPhen 2018 in Lyon.

## Author contributions

Conceptualization: AF, VD and RC. Flies collection: AF and VD. Experimentation and microscopy data collection: AF. Photogrammetric data collection: CVG and AD. Morphometric analysis: CVG. Results, discussion and writing: CVG, VD and RC.

## Data accessibility

All data and materials are available at the Dryad Digital Repository.

